# Effects of a novel proteasome inhibitor, UR238 on the tumor immune microenvironment and growth in epithelial ovarian cancer

**DOI:** 10.1101/2025.11.21.689729

**Authors:** John Phillip Miller, Naixin Zhang, Kyukwang Kim, Cameron WA Snyder, Jamie Linn McDowell, Michelle Elizabeth Whittum, Negar Khazan, Niloy Singh, Myla Strawderman, Richard Moore, Rakesh Singh, Rachael B. Rowswell-Turner

## Abstract

High-grade serous ovarian cancer (HGSOC) is a deadly gynecologic malignancy, often diagnosed at an advanced stage and most patients will experience recurrence and resistance to platinum-based chemotherapy. While there are few targeted therapies available for HGSOC, there are no effective immunotherapies available to treat this disease.

The ubiquitin-proteasome system (UPS) maintains cellular protein homeostasis by degrading misfolded or damaged proteins. In epithelial ovarian cancer (EOC) elevated expression of proteasome subunit PSMB4 correlates with epithelial ovarian cancer growth and poor prognosis. A proteasome inhibitor has yet to be approved for EOC treatment despite evidence of activity in phase-1/2 clinical trials. Limitations of past generation proteosome inhibitors include sub-optimal solid tumor/tissue penetration due to boronic acid functionality, and poor solubility.

Here, we show that a novel proteasome inhibitor, UR238, suppresses viability of several EOC cell lines and reduces tumor burden in both murine xenograft and rat syngeneic models. In a rat EOC model, UR238 treatment reduced tumor burden, shifted the predominantly suppressive immune microenvironment to an inflammatory phenotype with reduced suppressive macrophages, and decreased PD-L1 expression on myeloid cells. Additionally, in a mouse model, UR238 also induced robust immune changes in HGSOC bearing mice without a corresponding reduction in tumor burden. Our data highlights the promise of UR238 for the treatment of ovarian cancers as a single-agent or in combination with immunotherapies, where modulating the immune composition of these tumors will be important in improving outcomes for ovarian cancer patients.

## Introduction

In the U.S., ovarian cancer is the second most common malignancy of the gynecologic tract, yet it is the most deadly ^1^. High-grade serous ovarian cancer (HGSOC) is the most common histologic subtype, accounting for over 60% of all ovarian malignancies ^2^. Although commonly referred to as “ovarian” cancer, recent studies suggest that HGSOC originates from the fallopian tube epithelium ^3^. HGSOC is typically diagnosed at advanced stages due to vague symptoms such as bloating, fatigue, or abdominal pain, as well as the lack of an effective screening test ^4^. The current treatment paradigm for newly diagnosed ovarian cancer is a combination of systemic therapy and surgical resection. First line systemic therapy for HGSOC is platinum-based cytotoxic chemotherapy, and this has been the mainstay of treatment for decades. In more recent years the benefit of targeted therapies, such as PARPis, have been realized for ovarian cancers with homologous recombination deficiency. Unfortunately, HGSOC often recurs after initial treatment and once it does, cure is no longer possible, and treatment becomes palliative in nature. Furthermore, the treatment options become increasingly limited at each subsequent recurrence. This highlights an unmet need for more therapeutic options in the population of heavily pretreated HGSOC patients.

The immune system is important in both promoting ovarian cancer development, And controlling its progression ^5,6^. Specifically, Zhang et al. found that tumoral T cell infiltration correlates with improved survival for ovarian cancer patients. On the other hand, suppressive cells such as regulatory CD4^+^ T cells and tumor-associated macrophages (TAMs) can dampen the antitumor immune response ^7^. Immunotherapy is an emerging type of systemic therapy that utilizes the body’s own immune system to target cancer cells. The most heavily utilized immunotherapies in oncology today are checkpoint inhibitors, which act by blocking checkpoint ligands, such as PD-1 and PD-L1. These new treatments, when used alone or in combination with cytotoxic chemotherapy, have resulted in increased response rates and response durations across many different cancers. Immunotherapy typically has a more tolerable side effect profile when compared to cytotoxic chemotherapy. In gynecologic oncology, immunotherapies have been FDA approved in endometrial and cervical cancer, but the efficacy of immunotherapy has been limited in ovarian cancer clinical trials thus far. One potential reason for this is the abundance of TAMs, a major driver of tumor progression and immune suppression in the ovarian TME and associated malignant ascites. Although macrophages are classically divided into anti-tumor M1 and tumor-promoting M2 to describe their phenotype, they can be plastic and exist along a spectrum of function and phenotype. The depletion, inhibition or repolarization of suppressive macrophages represents a promising putative immunotherapy strategy in ovarian cancer ^8^.

The ubiquitin-proteasome system (UPS) involves ubiquitination and proteolysis of proteins by a series of enzymes. The UPS is essential in maintaining the stability of proteins that regulate cell cycle, DNA repair and transcription, and inhibition of this process can ultimately impact numerous cellular processes in cancer cells and immune cells ^9,10^. Proteasome inhibitors are a class of drugs that block this proteasomal protein degradation, leading to a build-up of ubiquitinated proteins and ultimately, cell cycle arrest, and apoptosis. This process occurs in both cancer cells and suppressive TAMs, making this an appealing target for cancer therapies ^9–12^. It is not entirely clear how proteosome inhibitors impact the TME, but proteasomal degradation of I-kB has been shown to regulate NF-kB activity by allowing NF-kB to translocate to the nucleus. Therefore, proteasomal inhibition leads to increased I-kB, and therefore NF-kB is sequestered in the cytosol. The NF-kB pathway controls many processes that are exploited by cancer cells to facilitate their growth. Therefore, proteasome inhibition leading to NF-kB downregulation is a potential strategy for reversing immune evasion in ovarian cancer ^14,15^.

Proteosome inhibitors are currently an effective FDA-approved treatment for multiple myeloma and lymphoma. However, their benefit in ovarian cancer is undefined. Although proteasome inhibitors commonly used in liquid cancers, phase I trials have shown some impact in ovarian cancer ^16,17^. Bortezomib was the first proteasome inhibitor to be FDA-approved in 2005 as a multiple myeloma therapy ^10,14,18^. Carfilzomib was the second-in-class proteasome inhibitor drug to receive fast-track FDA approval in 2012 for relapsed and refractory multiple myeloma ^10^. Carfilzomib irreversibly binds to and is more selective to the proteasome and therefore has improved safety profiles compared to bortezomib and more durable inhibition ^19^. However, carfilzomib has poor aqueous solubility making it difficult for the drug to be absorbed, necessitating 50-fold excess of b-cyclodextrin derivative to prepare a solution to inject ^10^. Due to these limitations, new proteasome inhibitors as cancer therapies are an active area of development ^10,14,19^.

Here, we show how a novel proteasome inhibitor, UR238, functions to reduce ovarian cancer cell viability while regulating the tumor immune microenvironment in multiple human and animal models of ovarian cancer. Furthermore, we show that UR238 reduces levels of the immunomodulator HE4 *in vitro*, highlighting the drug as a potential therapeutic for HGSOC.

## Results

### UR238 functions as a proteasome inhibitor in vitro

Proteasome inhibitors have been shown to be effective anti-cancer therapies for liquid tumors, namely multiple myeloma ^20^, yet have thus far had limited efficacy in solid tumors. We previously developed a proteasome inhibitor with improved drug characteristics with the goal of increasing effectiveness in solid tumors by optimizing bioavailability and tissue penetrance to deliver sufficient doses of drug to tumor tissue, including gynecologic malignancies. We began with a carfilzomib backbone and chemically modified the structure to improve the calculated LogP (cLogP), which is a measure of a compound’s hydrophilicity and thus predicts enhanced absorption into tissue. Figure 1 A shows the structural modifications that resulted in the development of UR238 (cLogP of 5.09). To confirm proteasome inhibition at the cellular level, we treated OVCAR-3 cells, an ovarian cancer cell line, with varying doses of UR238; the total ubiquitinated proteins in the cell lysate increased, further supporting its function as a proteasome inhibitor in ovarian cancer cells (Fig. 1 B).

**Figure 1:**
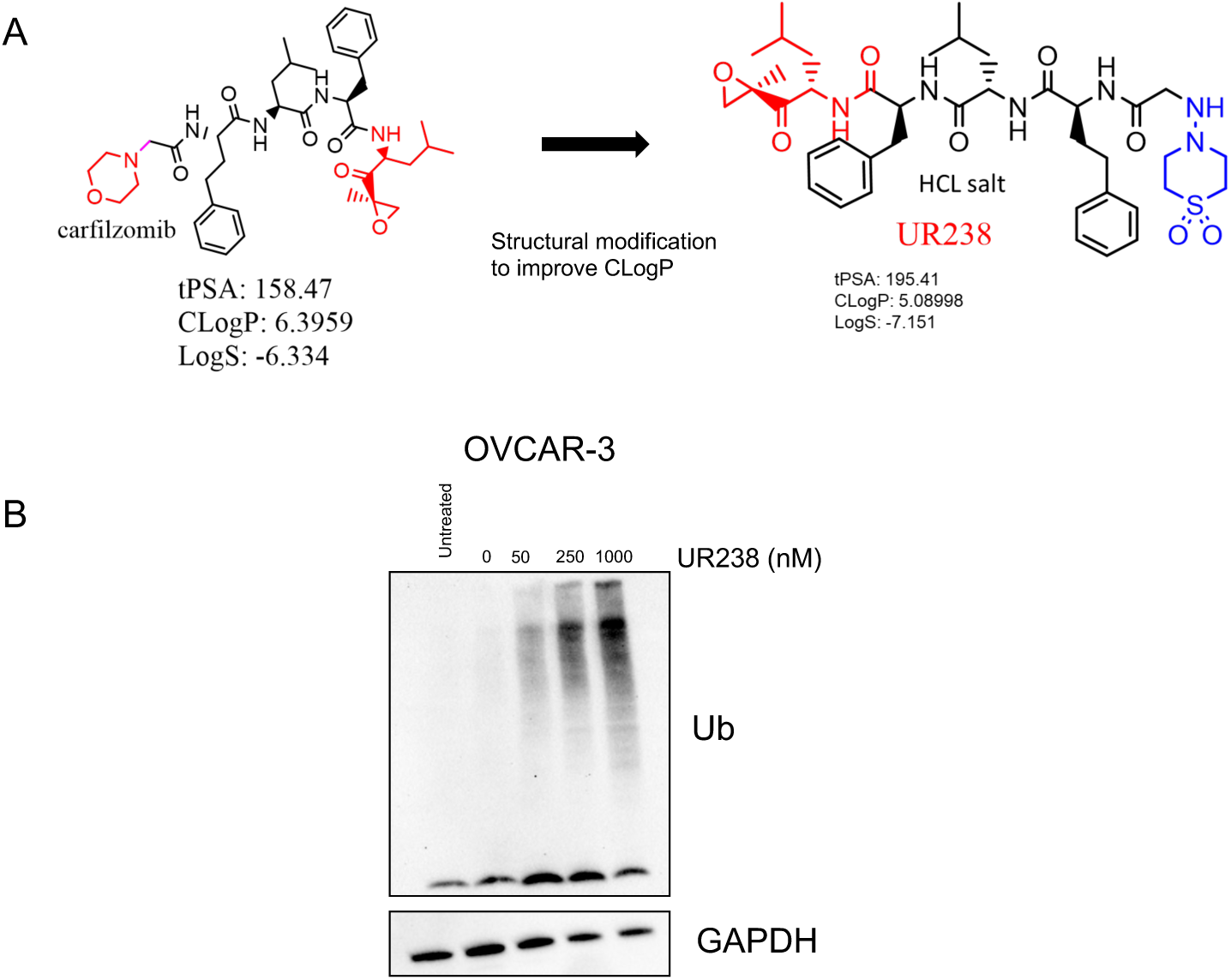
UR238 is an effective proteasome inhibitor. A) Chemical structure of carfilzomib and UR238 with respective tPSA, CLogP, and LogS values. B) OVCAR-3 cells were treated with UR238 at 0-1000 nM for 24 hours and total ubiquitin was analyzed by western blot analysis.

### UR238 decreases ovarian cancer cell viability in vitro and in vivo

Next, we wanted to determine how UR238 impacts the viability of cancer cells at various doses and timepoints. We treated a panel of human ovarian cancer cells (OVCAR-3, OVCAR-8, SKOV3) with UR238 for 24, 48 and 72 hours and assessed their viability using an MTS assay. After 48 hours, there was a decrease in cell viability at concentrations of 1μM or higher (Fig. 2 A). There was also a decrease in viability at less than 0.25 μM after 72 hours.

**Figure 2:**
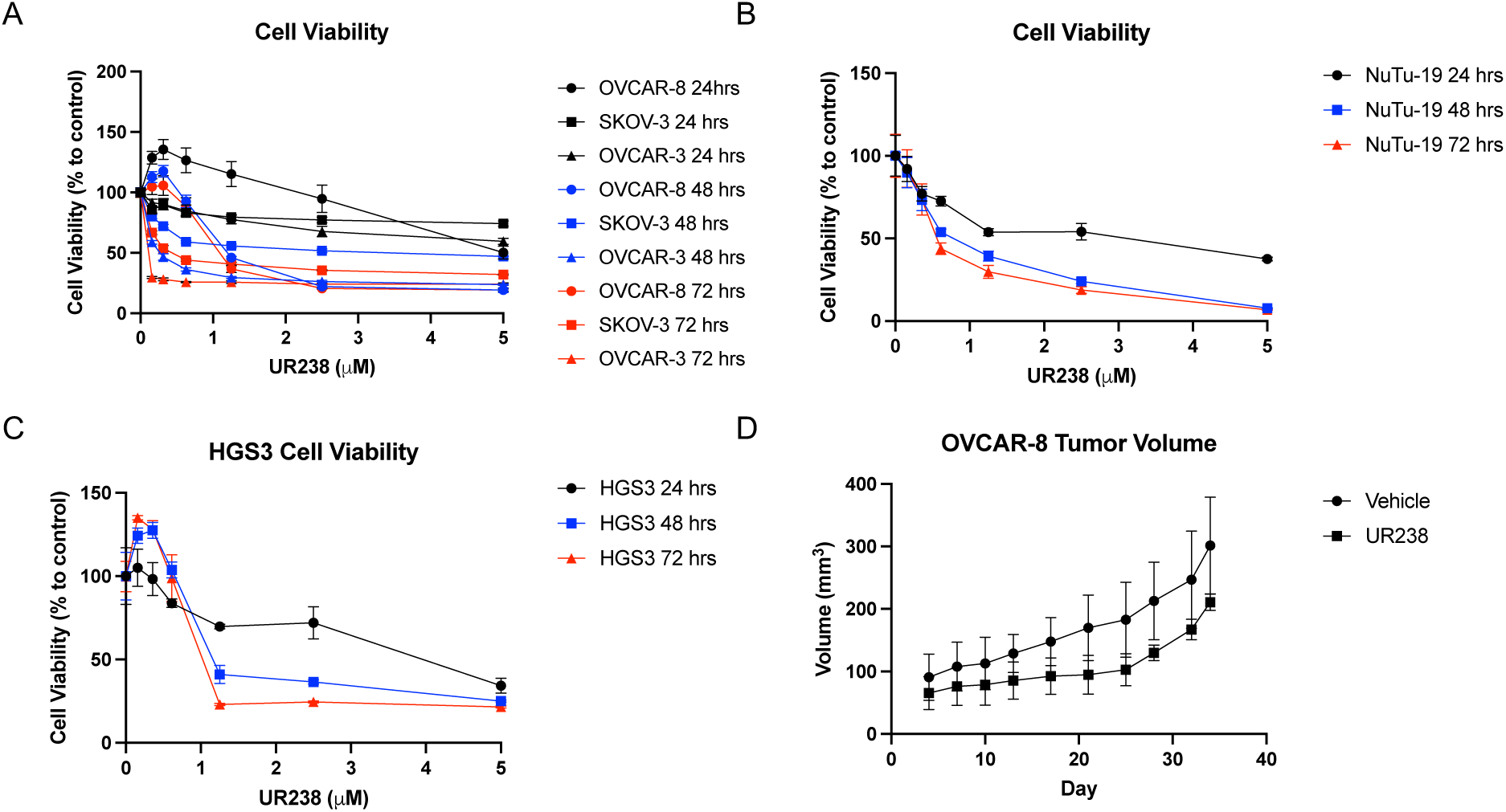
UR238 decreases ovarian cancer cell viability in vitro and in vivo. A) Cell viability of OVCAR-8, SKOV-3, OVCAR-3 cells after treatment with UR238 for 24, 48 and 72 hrs was assessed by MTS assay. B) Cell viability of NuTu-19 cells after treatment with UR238 for 24, 48, and 72 hrs was assessed by MTS assay. C) Cell viability of HGS3 cells after treatment with UR238 for 24, 48, and 72 hrs was assessed by MTS assay. D) OVCAR-8 cells were implanted subcutaneously into NSG mice and tumor growth was measured over time with UR238 (2mg/kg, MWF) or vehicle treatment starting at day 9 post-injection. T-test was used to determine there was a statistically significant difference (p≤0.05) in tumor volume on days 14-32.

UR238’s impact on ovarian cancer cells also translates to two rodent cell lines, the NuTu19 rat ovarian cancer line and the HGS3 fallopian tube murine cancer model. We tested *in vitro* viability in the rat ovarian cancer cells following UR238 treatment and found NuTu19 exhibited similar decreases in viability after 48 to 72 hours (Fig. 2 B).

The murine HGS3 cell line was generated by the Drapkin laboratory from a spontaneous tumor developed using the Cre-LoxP system to induce deletion of tumor suppressor genes specifically in the fallopian tube ^21^. The fallopian tube has recently been recognized as the primary cell of origin for human HGSOC ^22^.

Using a murine tumor cell line of fallopian tube origin that mimics the genetic abnormalities of human tumor cells helps ensure a clinically relevant murine ovarian cancer model. We again treated these HGS3 cells with varying doses of UR238 for 24, 48 and 72 hours and assessed their viability using an MTS assay. Similarly to the human cell lines, HGS3 cell viability decreased at doses above 1μM after 48 and 72 hours (Fig. 2 C).

To test UR238’s effectiveness on cancer cells *in vivo*, first, a human xenograft model was employed, as this allows for the most efficient pre-clinical drug modeling experiments. We injected NSG mice with 1 x 10^6^ OVCAR-8 cells subcutaneously and treated with UR238 2 mg/kg, 2-3 times per week for 34 days. Figure 2 D shows the tumor volume measured over time. We observed a significant difference in tumor volume between control and treated mice at days 14-32 (Fig. 2 D). The xenograft model, however, did not allow us to assess the impact of UR238 on the tumor immune microenvironment. Therefore, we utilized an immune competent rat model of ovarian cancer, NuTu19, to assess the impact of UR238 on tumor burden and the immune microenvironment.

### UR238 reduces tumor burden in a rat syngeneic model of ovarian cancer

To determine the effect of UR238 on the immune microenvironment of ovarian cancer, the NuTu19 cell line was injected into the peritoneal cavity to model metastatic carcinomatosis. Once injected, these cells seed the omentum, the most common site of ovarian cancer metastases in human patients ^23^. We treated the animals with either vehicle or UR238 (2 mg/kg) three times per week for 60 days. Rats treated with UR238 showed a significant reduction in ascites volume and omental tumor mass (Fig. 3 A, B) indicating a reduction in tumor burden and disease. The immune cells of the omentum and ascites were also analyzed by flow cytometry. UR238 treated rats had a reduction in macrophages in both the omentum and the ascites (Fig. 3 C, D). Macrophages in the ascites also had reduced expression of PD-L1 (Fig. 3 C, Right). UR238 also reduced the percentage of granulocytes in the omentum (Fig. 3 E, left). Additionally, UR238 treated rats also had a reduced percentage of granulocytes expressing PD-L1 in the omentum (Fig. 3 E, right). We also performed a Luminex analysis to determine the cytokine profile of the omentum and ascites of the rats to better understand the functional differences in the tumor and peritoneal cavity after UR238 treatment. We found that IL-10, an immunosuppressive cytokine, was decreased in the omentum and ascites of UR238 treated rats (Fig. 3 F). Additionally, we observed a decrease in the chemokine CCL2 in the omentum and the ascites, which is important in the trafficking of macrophages from circulation into tissues such as the tumor (Fig. 3 F). Together, this suggests that UR238 may cause a shift in the peritoneal cavity and tumor microenvironment away from a more immunosuppressive environment characterized by the decrease in the suppressive cytokine IL-10, and a decrease in the macrophage attractant CCL2, leading to reduced percentages of macrophages in the peritoneal cavity and tumor bearing omentum.

**Figure 3:**
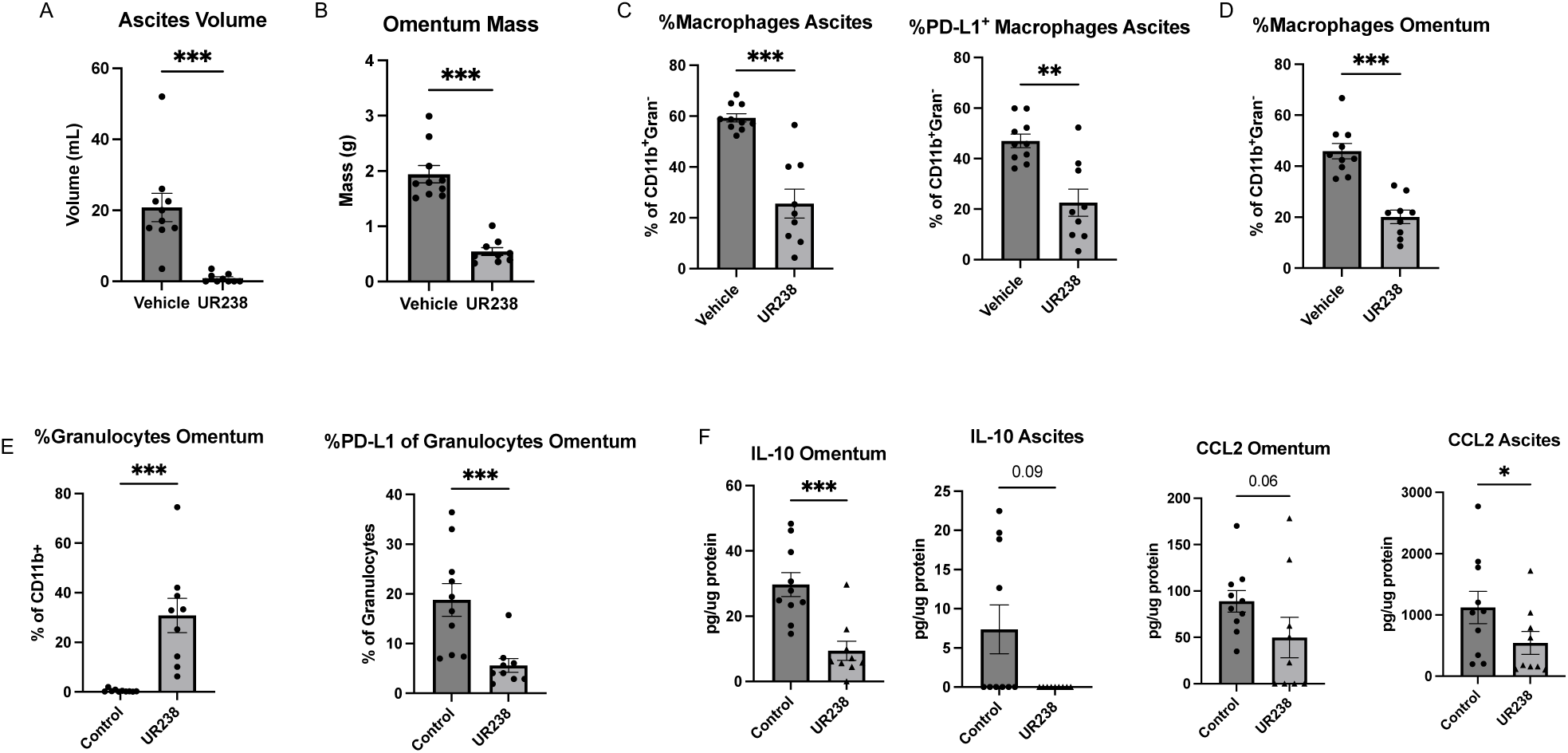
UR238 reduces tumor burden in a rat syngeneic model of ovarian cancer. Rats were injected intraperitoneally (i.p) with NuTu19 cells and treated with vehicle or UR238 (2mg/kg) i.p. and sacrificed 60 days later. A) Bar graph showing ascites volume collected and measured from vehicle and UR238 treated rats. B) Bar graph showing the tumor bearing omentum mass between vehicle and UR238 treated rats. C) The percentage of macrophages (CD11b+CD3e-CD45+MHCII+MacSub+) (left) and macrophages expressing PD-L1 (right) in ascites of vehicle or UR238 treated tumor bearing rats. D) The percentage macrophages in tumor bearing omentum of vehicle or UR238 treated tumor bearing rats. E) The percentage of granulocytes (CD11b+CD3e-CD45+Gran+) (left) and granulocytes expressing PD-L1 (right) in the omentum between Vehicle and UR238 treated rats. F) IL-10 and CCL2 levels were determined using a Luminex assay using samples from the omentum and ascites comparing Vehicle and UR238 treated rats.

### UR238 decreases PDL1 expression on myeloid cells in human ascites samples

To test the direct effect of UR238 on the tumor immune microenvironment in human patient samples, we used freshly isolated HGSOC tissue, and ascites, processed into single cell suspensions. This suspension contained both cancer cells and immune cells and was treated with UR238 or vehicle control for 48 hours. We then processed the cells for flow cytometry to identify immune cell population changes *in vitro* after drug treatment. We found that UR238 treatment led to a decrease in PD-L1 expression on CD11b^+^ myeloid cells in the ascites from these patients (Fig. 4 A). However, when the tumor cultures were treated with UR238, the same change in PD-L1 expression was not observed (Fig. 4 C). This could indicate that the myeloid cells from the tumor microenvironment are in a more polarized, less plastic state, where their immunosuppressive phenotype cannot be altered by UR238 alone.

**Figure 4:**
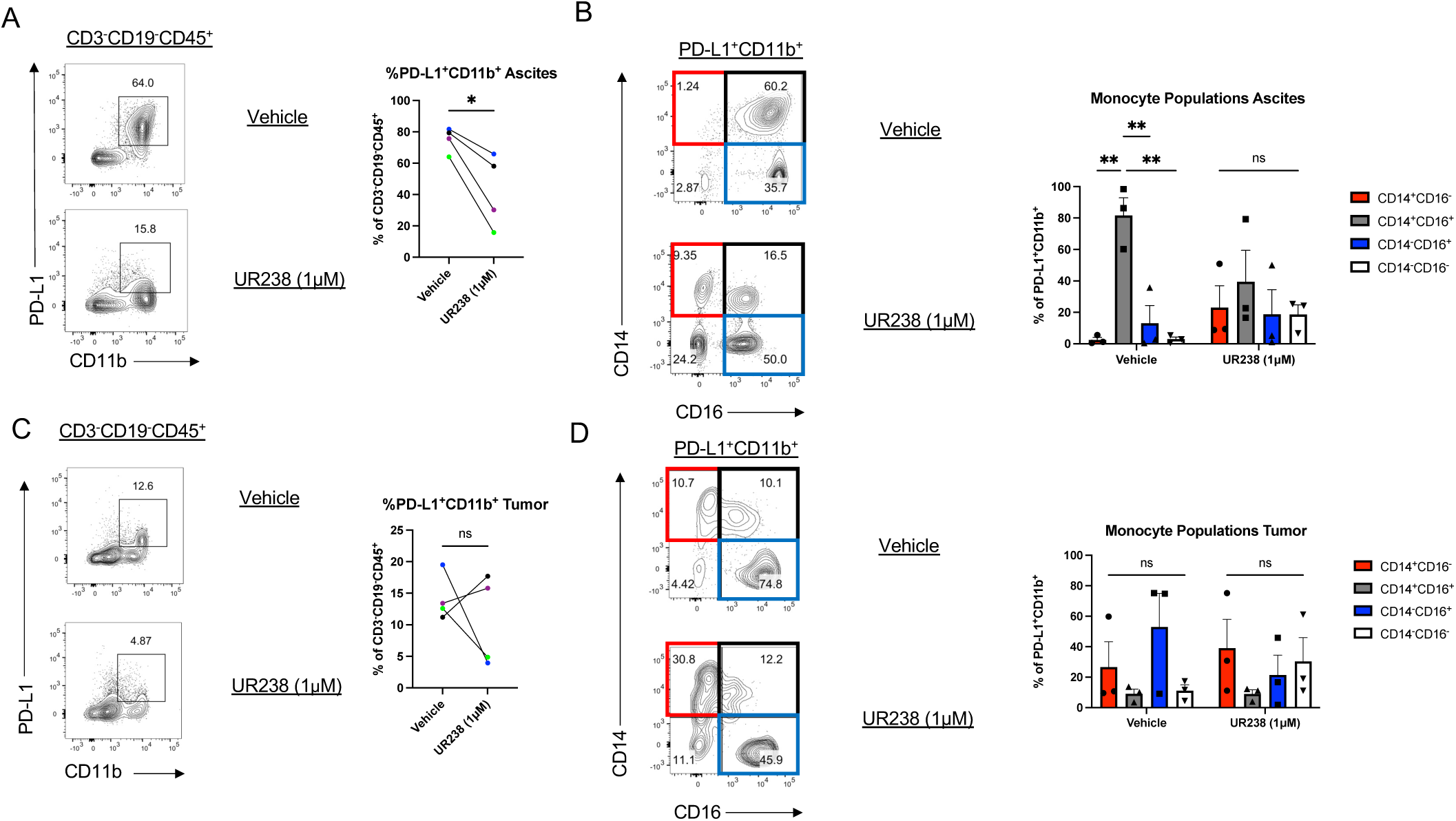
UR238 decreases PDL1 expression on myeloid cells in human patient ascites samples, but not tumor samples. Human tumor and ascites samples were processed to single cell suspensions, plated, and treated with vehicle or UR238 for 48 hours. A) Flow cytometry plots showing PDL1 and CD11b expression on CD3^-^CD19^-^CD45^+^ cells from ascites cultures. Percentage of PD-L1^+^CD11b^+^ cells is shown to the right. Paired t-test, *p < 0.05. B) Flow cytometry plots showing CD14 and CD16 expression among the gated PD-L1^+^CD11b^+^ cells in A. The graph to the right shows the percentages of the indicated monocyte populations. C) Flow cytometry plots showing PDL1 and CD11b expression on CD3^-^CD19^-^CD45^+^ cells from tumor cultures. D) Flow cytometry plots showing CD14 and CD16 expression among the gated PD-L1^+^CD11b^+^ cells in C. The graph to the right shows the percentages of the indicated monocyte populations. A and C used paired t-test. B and D used Two-Way ANOVA (ns p>0.05, *p<0.05, **p<0.01, ***p<0.001).

We further analyzed the myeloid cells within these PD-L1^+^CD11b^+^ cells by looking at additional phenotypic markers. Monocytes are mononuclear phagocytes that develop and differentiate from a common myeloid progenitor (CMP) in the bone marrow and then circulate in the blood, with the capacity to infiltrate tissues ^24^. Monocytes are a heterogenous cell population that can be separated based on cell surface markers into classical/inflammatory (CD14^+^CD16^-^), intermediate (CD14^+^CD16^+^) and non-classical/anti-inflammatory (CD14^-^CD16^+^) monocytes. We analyzed CD14 and CD16 expression among the PD-L1^+^CD11b^+^ myeloid cells to understand how these populations were changing with UR238 treatment. The proportion of CD14 and CD16 expressing monocytes shifted within the ascites after treatment, indicating a more inflammatory phenotype defined by more CD14^+^CD16^-^ cells, and less intermediate CD14^+^CD16^+^ monocytes as compared to control cultures (Fig. 4 B). However, within the tumor cultures, which already contained fewer intermediate monocytes, there was minimal change compared to the ascites (Fig. 4 D). Altogether, this data indicates that UR238 influences the immune microenvironment in human *in vitro* ascites cultures, with a shift towards a less immunosuppressive environment especially in the ascites, while tumor cultures remain more static.

### Mice bearing ovarian cancer peritoneal metastases have decreased M2 macrophages in the peritoneal cavity after UR238

To better understand the immune microenvironment in ovarian cancer, we began to utilize the HGS3 mouse model because of the superior availability of antibodies to characterize immune cell subsets in response to proteasome inhibition by UR238.

To validate the effect of UR238 on myeloid cells in the peritoneal cavity of tumor bearing mice, we injected HGS3 cells IP into female C57BL/6 mice ^21^. Mice developed peritoneal tumors that visibly seeded the omentum at 6 weeks post-injection. The peritoneal cavity contains a variety of leukocytes that patrol and migrate in and out of the omentum, where they encounter antigens or pathogens and mount immune responses ^25^. Immune cells in the peritoneal cavity typically function to preserve tissue homeostasis and facilitate tissue repair. As such, macrophages are a dominant cell type found in the peritoneal cavity. Following stimulation, monocytes can be recruited to the peritoneal cavity where they can polarize to an inflammatory (Ly6C^hi^) or anti-inflammatory (Ly6C^low^) phenotype ^26^.

Because the peritoneal cavity can be an important source of infiltrating immune cells into peritoneal tumors, we aimed to characterize the immune populations in HGS3-tumor bearing mice in the peritoneal cavity. Compared to non-tumor bearing mice, HGS3-bearing mice have an increase in the percentage of Ly6C^+^F4/80^+^ inflammatory macrophages, indicating that this population may be important in the tumor response in the peritoneal cavity (Fig. 5 A). We then treated tumor bearing mice with UR238 or vehicle control starting 1-week post-tumor cell inoculation and continued 3 times per week at 2 mg/kg, for 6 weeks. The tumors and peritoneal lavage were isolated to determine efficacy and immune changes *in vivo* following treatment. Although there was no significant difference in tumor mass between the groups (data not shown), there were significant shifts in the myeloid populations of the peritoneal cavity. Ly6C^+^F4/80^+^ macrophages were significantly increased compared to Ly6C^-^F4/80^+^ macrophages in the UR238 treated group (Fig. 5 B). We also cultured macrophages from the peritoneal cavity of untreated HGS3-tumor bearing mice and treated the cultured macrophages with UR238 or DMSO vehicle control for 48 hours and analyzed the cells by flow cytometry. We found that the myeloid cells treated with UR238 showed a dose dependent reduction in the expression of PD-L1 (Fig. 5 C).

**Figure 5:**
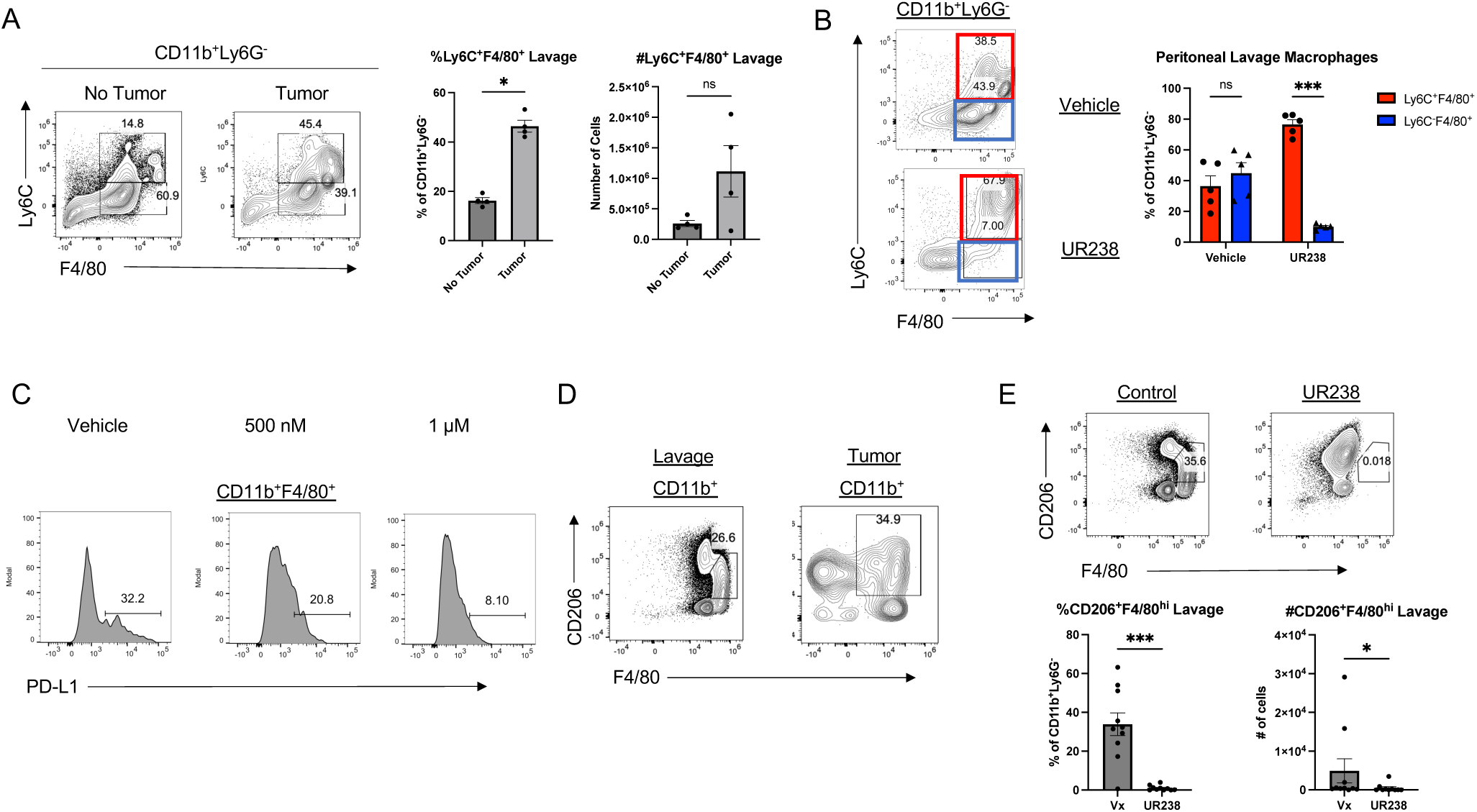
Mice bearing ovarian cancer peritoneal metastases have decreased M2-like macrophages in the peritoneal cavity after UR238 treatment. A) Flow cytometry plots showing the proportions of Ly6C^+^F4/80^+^ peritoneal macrophages in non-tumor bearing mice vs tumor bearing mice. Percent and cell number of Ly6C^+^F4/80^+^ macrophages are quantified to the right. B) Flow cytometry plots showing the percentage of Ly6C^+^F4/80^hi^ and Ly6C^-^F4/80^hi^ cells in the peritoneal. C) PD-L1 expression on ex vivo cultured peritoneal macrophages from a tumor bearing mouse that were treated with UR238 or control for 48 hours. D) Flow cytometry plots showing CD206 and F4/80 expression among CD11b^+^ myeloid cells in the peritoneal lavage and tumor. E) CD206 and F4/80 expression in peritoneal macrophages from vehicle (Control) vs UR238 treated mice. Percent and number of CD206^+^F4/80^+^ macrophages is quantified below. Mann-Whitney U Test (ns p>0.05, *p<0.05, **p<0.01, ***p<0.001).

Additionally, continuing our characterization of the immune cells in the peritoneal cavity and tumor, we found that untreated mice contain F4/80^hi^CD206^+^ macrophages in the peritoneal cavity and CD206^+^F4/80^+^ macrophages in the tumor (Fig. 5 D). CD206 expression is indicative of an M2 suppressive macrophage phenotype ^27^. When treated with UR238, we observed a significant reduction in the CD206^+^F4/80^hi^ population of macrophages (Fig. 5 E). This data suggests that UR238 is shifting the myeloid compartment of the peritoneal cavity specifically toward a more M1, pro-inflammatory phenotype based on the increase in Ly6C^+^ macrophages and the reduction of CD206^+^ macrophages. Interestingly, this M2 to M1 shift is not seen in the tumor of these mice, where the CD206^+^ macrophage population remains similar between control and UR238-treated mice (Fig. 6).

**Figure 6:**
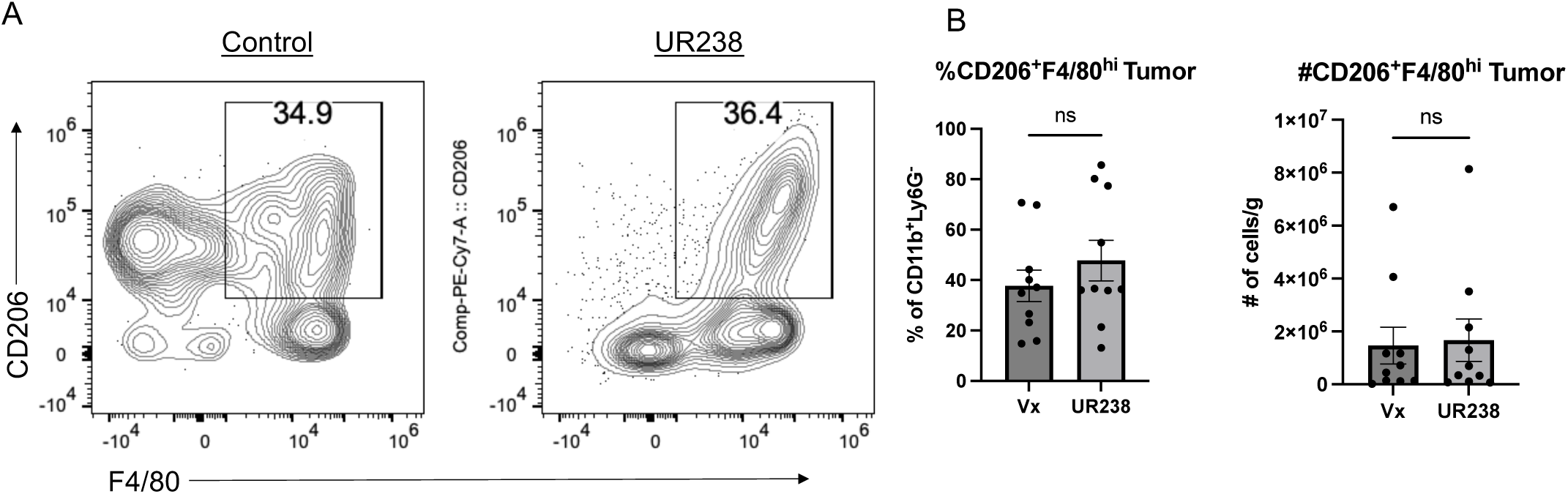
CD206^+^F4/80^hi^ macrophages remain unchanged in tumors following UR238 treatment. A) Flow cytometry plots showing F4/80 and CD206 expression in control and UR238 treated tumors. B) Percent and cell number per gram of tumor for the CD206+F4/80hi population from A. Mann-Whitney U Test (ns p>0.05, *p<0.05, **p<0.01, ***p<0.001).

**Figure 7:**
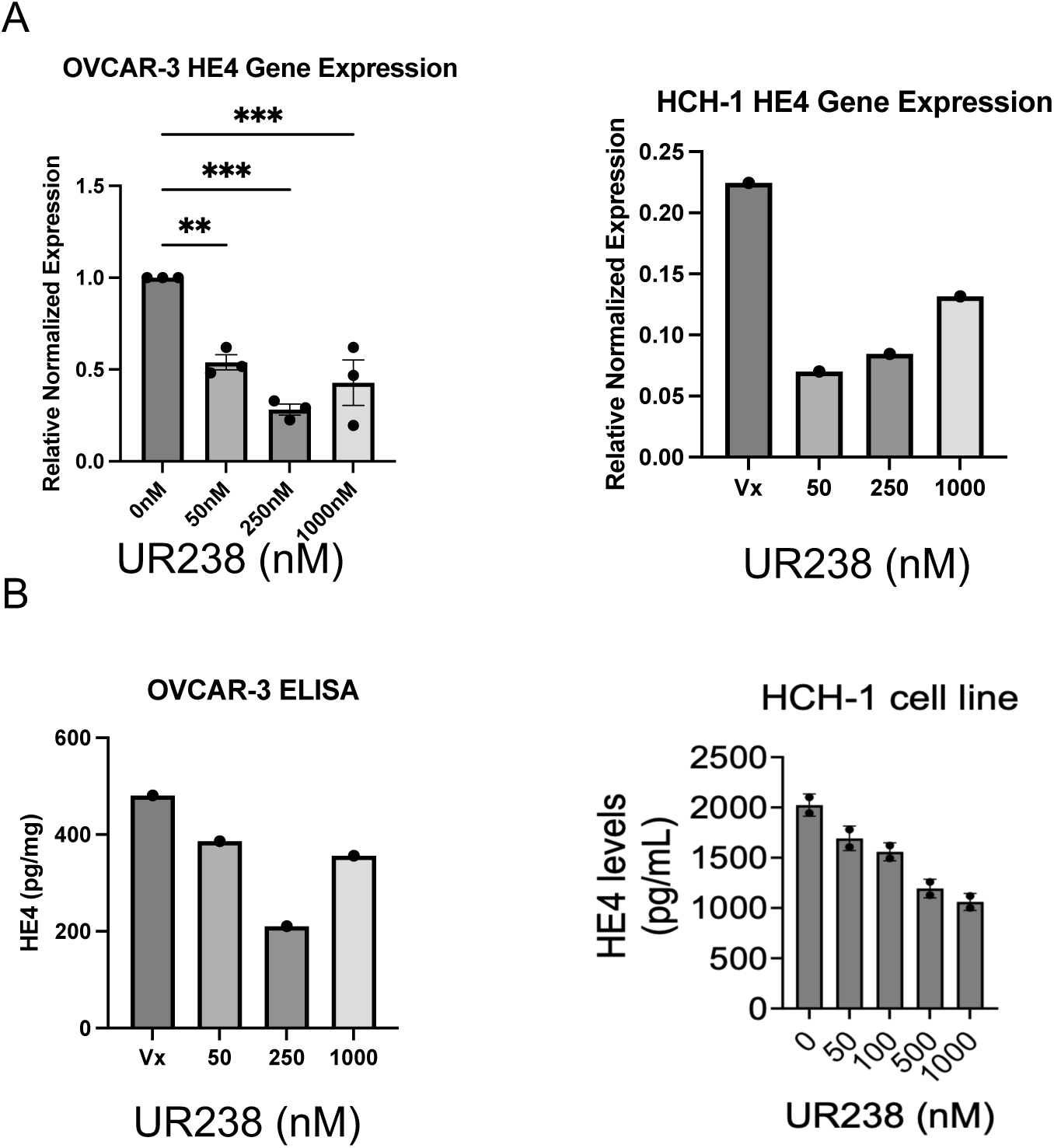
UR238 decreases HE4 protein expression in ovarian cancer cell lines in vitro. A) qPCR analysis of *WFDC2* expression in OVCAR-3 and HCH-1 cells. Each dot represents the mean of three technical replicates. B) HE4 expression by ELISA in OVCAR-3 and HCH-1 cell lysate treated with UR238 for 24hrs. Each dot represents the mean of two technical replicates. HCH-1 data represents technical replicates from one experiment. One-way ANOVA (ns p>0.05, *p<0.05, **p<0.01, ***p<0.001).

Since we identified changes in PD-L1 in the data shown above, we hypothesized that UR238 in combination with PD-L1 blocking antibodies would synergize to improve the tumor response. We treated HGS3-bearing mice with UR238 and anti-PD-L1 or isotype control antibodies three times per week. Surprisingly, we found no significant difference in tumor burden between the treatment groups (Fig. S1 A). Additionally, there was still a significant decrease in M2 macrophages in the peritoneal cavity in both UR238 treated groups, but not in the Anti-PD-L1 single treatment group (Fig. S1 B). Additionally, UR238 + anti-PD-L1 showed no significant difference in M2 macrophages within the tumor itself (Fig. S1 C). Overall, this highlights a need for more research into potentially other treatment schedules, dosages, or utilizing blockade of different inhibitory receptors.

### UR238 decreases HE4 protein expression in ovarian cancer cell lines in vitro

HE4 is a protein of the Whey Acidic Protein (WAP) family of immunomodulators that has been shown to function as a serum biomarker for ovarian cancer diagnosis ^28^. HE4 serum levels correlate with worse survival for patients with all stages of ovarian cancer ^29^. The FDA approved Risk of Ovarian Malignancy Algorithm (ROMA) uses HE4 and another biomarker, CA125, to classify patients into high or low risk groups with high accuracy ^30^. We have previously shown that HE4 overexpression in human HGSOC promotes a suppressive tumor immune microenvironment ^31^. Therefore, HE4 is an attractive target for treatments that lead to decreased HE4 production in HGSOC. To this end, we wanted to determine if UR238 had any impact on HE4 levels in the human cell lines available in our lab. To determine if UR238 was reducing HE4 at both the gene and protein level, we treated two cell lines with varying doses of UR238 and assessed gene and protein expression by RT-qPCR and ELISA (Fig. 8 A, B). It was found that UR238 significantly decreased transcription of *WFDC2* in OVCAR-3 and HCH-1 cells and reduced protein expression in cell lysates by ELISA. More work needs to be done to determine the mechanism by which UR238 is leading to a reduction in HE4 levels, but this data indicates that UR238 treatment may modulate the immune microenvironment as well as reduce a known ovarian tumor biomarker.

## Discussion

Proteasome inhibition is a successful treatment strategy in liquid cancers, such as multiple myeloma, but this class of drugs have yet to show clear benefit for patients with solid malignancies ^32,33^. High expression of PSMB4 has been shown to correlate with tumor growth and poor prognosis in ovarian cancer patients ^34^. This study explores the therapeutic potential of a novel proteasome inhibitor in ovarian cancer.

Two main chemical classes of proteasome inhibitors have been used clinically: boronic acid (bortezomib, ixazomib) and epoxymycin (carfilzomib). Bortezomib, a first-in-class 26S proteasome inhibitor approved for hematologic malignancies, has been shown to effectively sensitize EOC cells to chemotherapeutics, similar to paclitaxel and cisplatin in experimental models ^35^. However, it has only shown modest activity in phase-I and -II ovarian cancer clinical trials, even when combined with chemotherapy ^17,36^ We hypothesize that the chemical structures of both bortezomib (boronic acid) and poor cLogP of carfilzomib have hindered a successful solid cancer proteosome inhibitor clinical trial.

We therefore chose to modify carfilzomib to optimize its solubility following Lipinski’s drug-likeness rules, which predict that compounds with an optimal clogP (0-5) will be more likely to be a successful drug in the clinic. We replaced the morpholine moiety of carfilzomib with N-amino dioxothiomorpholine thus imparting a superior cLogP of 5.09.

Our findings demonstrate that our novel proteasome inhibitor, UR238, reduced ovarian cancer cell viability *in vitro* in multiple cell lines and furthermore, we also found that UR238 inhibited ovarian tumor growth *in vivo* in a rat syngeneic model, and a human tumor xenograft model. While this evidence supports that proteasome inhibitors have a direct effect on cancer cells, the promise of proteasome inhibitors lie in that their effect leads to tumor cell death as well as immune modulation. In line with a previous study by Zhou et al in which carfilzomib reprogrammed macrophages from an M1 to an M2-like phenotype, our data further supports that proteasome inhibitors may have an immune modulating effect, potentially enhancing cytotoxicity within the tumor microenvironment ^9^. Further research into how this class of drugs impacts the solid tumor microenvironment will be beneficial in developing new treatments for these patients.

Gynecologic malignancies develop and spread in a unique tumor microenvironment characterized by a predominance of suppressive macrophages ^37^. We demonstrated here that macrophages within the peritoneal cavity and tumor were significantly reduced in UR238-treated animals, indicating a shift towards a pro-inflammatory microenvironment. There was also a corresponding shift in the suppressive cytokine IL-10 and the monocyte/macrophage chemoattractant CCL2 in the tumor and ascites of treated rats. Thus, our work provides insight into the impact of proteasome inhibitors on the solid tumor microenvironment and surrounding cells within the peritoneal cavity ^9^. The question remains if UR238 will be effective as a monotherapy for solid tumors. While we showed *in vivo* efficacy in syngeneic rat and human xenograft models, we did not see a reduction in the HGS3 mouse model of ovarian cancer. This is a relatively new, but clinically relevant model of fallopian tube-origin EOC, which may be less immunogenic at baseline ^3^. Therefore, it is likely a better representation of the unique immune microenvironment in these classically immunotherapy-resistant cancers. Although UR238 was ineffective at reducing tumor-burden in HGS3-bearing mice, the shifts in immune phenotype can indicate a more favorable environment for treatment with immunotherapy such as checkpoint inhibitors.

Interestingly, based on our *in vitro* human tumor culture data, immune cells from tumors treated in culture with UR238 seem to be less plastic than cells isolated from the ascites of the same patients. This indicates that UR238 alone may not be enough to re-shape the immune microenvironment within the tumor and drive a therapeutic response. Thus, it is likely that for UR238 to be an effective immunotherapy agent for ovarian cancer patients in the future, it would need to be combined with other immune modifying agents. Although we did not observe significant synergy with anti-PD-L1 blockade, further work needs to be done to determine if UR238 will synergize with other immunotherapies such as anti-CTLA-4 therapies. We hypothesize that this might be the case, since UR238 appears to be promoting a more inflammatory environment within the peritoneal cavity, which would surround peritoneal metastases and might provide a source of anti-tumor immune cells to infiltrate the tumor. However, since these macrophages are highly plastic, whether UR238 is fully converting M2 macrophages to M1 macrophages remains to be determined ^38^. More functional assays determining whether UR238 treated macrophages suppress T cell function or increase phagocytic activity will be important next steps in understanding their role in the tumor microenvironment. An important aspect in future experiments and treatments would also be to determine if these peritoneal cavity immune myeloid cells are indeed infiltrating the tumor and retaining their inflammatory phenotype or immediately becoming suppressive TAMs upon infiltration.

We have also found that UR238 decreases gene and protein expression of the ovarian cancer biomarker HE4 *in vitro.* HE4 is a part of the WAP domain family of proteins, which are known anti-inflammatory effector proteins ^39^. HE4 serum monitoring is used in ovarian cancer clinical diagnosis and surveillance. Higher levels of this protein are associated with a poor prognosis, enhanced macrophage infiltration, reduced CD8^+^ T cell infiltration, and higher PD-L1 expression in patients. In animal models, forced HE4 over-expression leads to higher tumor burden, and increased levels of CCL2, IL-10, along with suppressive macrophage infiltration ^31^. Although the specific function of HE4 is still unknown, it clearly functions as a promoter of tumor immune suppression in ovarian cancer, and inhibition of HE4 is a cancer treatment strategy that is currently under study in our lab. Interestingly, our characterization of the changes to the tumor microenvironment with UR238 treatment follows a reverse pattern to that seen with high HE4 levels in ovarian cancer. Previous work from our lab showed that the NF-kB pathway regulates HE4 expression ^40^. Indeed, the proteasome plays an important role in regulating the NF-kB pathway through the ubiquitin-mediated degradation of I-kB, which keeps the NF-kB complex sequestered in the cytosol ^41^. Therefore, we hypothesize that UR238 may impact HE4 expression through proteasome inhibition resulting in decreased NF-kB signaling. Future work will focus on addressing this hypothesis and elucidating whether UR238-mediated immune changes are a result of the drug acting on immune cells directly, or a result of the effects on the tumor cells themselves.

In conclusion, we present UR238 as a novel proteasome inhibitor that decreases ovarian cancer viability and growth *in vitro* and *in vivo*. It is also capable of shifting the immune cells within the peritoneal cavity of tumor bearing mice towards a more pro-inflammatory phenotype. Future work will involve elucidating the precise mechanisms of UR238-mediated anti-tumoral effects. The results described here provide evidence that UR238 may have a role in the development of new and effective combination immunotherapy for ovarian cancer.

## Materials and Methods

### Human patient consent and samples

Participants underwent informed consent prior to surgery for removal of high grade serous ovarian tumors. Tissue was collected per trial UGYO18036 at the University of Rochester Wilmot Cancer Institute, which completed extensive ethical review and IRB approval. Following surgical resection, a portion of the tumor and ascites was collected and placed into a sterile container, maintained on ice, and then transported immediately to the lab for processing. The solid tumor specimen was mechanically digested through a 50-gauge stainless steel wire mesh in a large petri dish. The tumor and ascites suspension were then centrifuged (300 x g, 5 mins, 4°C) and the red blood cells (RBCs) were lysed. The tumor and ascites suspensions were then plated and treated with various concentrations of UR238 or DMSO vehicle control. After 24-48 hours, adherent cells were isolated using TrypLE Express (Cat #: 12604013) and stained according to the flow cytometry protocol.

### Cell culture and MTS

OVCAR-3 (DMEM), SKOV3 (DMEM), OVCAR-8 (DMEM), NuTu-19 (RPMI), and HGS3 (DMEM) cells were cultured in their respective media supplemented with 10% FBS and 1% Pen-Strep. To detach cells, 0.05% Trypsin-EDTA was added to the plate and incubated at 37°C for 2-5 minutes. Cells were then collected in their respective medium and centrifuged at 300 x g for 5 minutes at 20°C. The cells were then counted using a hemocytometer and subsequently used for the desired application.

For MTS assays, the Promega CellTiter96 Aqueous One Solution Cell Proliferation Assay kit (Cat # G3582) was used according to the manufacturer’s protocol. Briefly, 5,000 cells were plated in 96 well plates for each time point (24 hours, 48 hours, 72 hours) in 100 μl and allowed to adhere for 24 hours. The cells were then treated with UR238 at the indicated doses by removing media and adding fresh media containing the drug. After treatment, CellTiter 96 Aqueous One Solution Reagent was added to the wells and incubated at 37°C for 1-4 hours in 5% CO_2_ atmosphere. The absorbance was read at 490 nm using a 96-well plate reader.

### Animals and tumor experiments

All experiments were approved by the University Committee on Animal Resources and all animals were maintained in a specific-pathogen free environment in the University of Rochester animal facility. 6–10-week-old Female C57BL/6 were implanted with ∼8 x 10^6^ HGS3 cells intraperitoneally and mice were sacrificed after 6-8 weeks. For UR238 treatment, the DMSO containing drug was dissolved in a vehicle containing 40% hydroxypropyl-beta-cyclodextran (HBC) and 5% Kolliphor HS15 (Sigma # 42966). Mice were injected intraperitoneally one week post tumor injection with the drug (2mg/kg body weight) or control three times per week. For anti-PD-L1 experiments, 200ug per mouse of either anti-PD-L1 antibody (BioXCell, BE0101, Clone: 10F.9G2) or IgG2b isotype control (BioXCell, BE0090, Clone: LTF-2) were injected i.p twice per week starting one week post-tumor cell injection.

NuTu19 cells were implanted into 7-9-week-old female Fischer rats as previously described ^31^. Rats were treated with UR238 (2 mg/kg) or control three times per week and were sacrificed after 8 weeks. The tissue from tumor bearing rats was processed as previously described ^31^ and tumor mass and ascites volume were collected.

For human xenograft experiments, 1 x 10^6^ OVCAR-8 cells were implanted subcutaneously into NSG mice. Once tumors were detectable at day 4 post-injection, UR238 or vehicle control injections were started at 2 mg/kg two to three times per week.

### Peritoneal macrophage isolation

Peritoneal macrophages were isolated by making a small incision on the skin and injecting 5 mL of PBS with a 20 G needle into the peritoneal cavity, massaging the abdomen for 10-15 seconds. Fluid was then withdrawn and the needle removed. The peritoneal fluid was dispensed into a tube on ice. Peritoneal cells were centrifuged at 300 x g for 5 mins at 4℃. Cells were resuspended in RPMI (10% FBS, 1% Pen/Strep) depending on the application.

### Flow cytometry

Omentum and/or tumors from HGS3 injected C57Bl/6 mice were minced in plain RPMI and incubated in collagenase (2 mg/mL), DNase (25 ug/mL), and hyaluronidase (0.1 mg/mL) for 30min-1hr at 37°C. The suspension was strained through 70 µm strainers, diluted with 10 mL of cold complete RPMI (10% FBS) and centrifuged at 300 x g for 5 mins at 4°C. The cells were resuspended in PBS supplemented with 5% FBS (FACS buffer) for cell counting.

The spleen was isolated and crushed through a 70 µm strainer in cold complete RPMI (10% FBS) and transferred to tubes and the cells were centrifuged at 300 x g for 5 mins at 4°C. The resulting pellet was resuspended in 1X Red blood cell lysis buffer for 4-5mins at room temperature with occasional shaking. The suspension was diluted with complete RPMI (10% FBS) and centrifuged at 300 x g for 5 mins at 4°C. The pellet was resuspended in PBS for cell counting.

The peritoneal lavage was collected using a 5ml syringe and 20G needle to inject 5ml of PBS into the peritoneal cavity. The abdomen was massaged for 10-15 seconds, and the peritoneal lavage was collected using the needle. The needle was removed, and the resulting suspension dispersed into a tube on ice. The suspension was centrifuged at 300 x g for 5 mins at 4°C and resuspended in 10ml of PBS and passed through a 70 mm strainer. The suspension was centrifuged at 300 x g for 5 mins at 4°C and resuspended in PBS for cell counting. All samples were stained with LiveDead Yellow dye (1 µL per 1 x 10^6^ cells), washed with PBS without FBS, and subsequently stained with the appropriate antibodies in FACS buffer. All samples were stained with antibodies at a dilution of 1:200 for flow cytometry for 30-60 minutes. The following antibodies were used: anti-CD8⍺ (364-0081-82), anti-CD3ε (740283), anti-PD-1 (752299), anti-CD4 (15-0041-81), anti-Ly6C (128005), anti-CD11b (560458), anti-CD19 (568287), anti-CCR2 (750042), anti-CD45 (563053), anti-CD206 (25-2069-42), anti-PD-L1 (558091), anti-Ly6G (108470), anti-CD11c (561241), anti-CD45RA (551402), anti-CD161a (565413), anti-CD45 (561586), anti-CD3 (563949), anti-CD4 (561833), anti-CD8⍺ (740139), anti-RP1 (554907), anti-RT1b (744127), anti-HIS36/MacSub (554901), anti-CD11b/c (562222), anti-CD4 (550057), anti-PD-L1 (PA5-20343). The BV421-anti-rabbit secondary antibody was used for anti-PD-L1 staining (565014). For intracellular staining, surface staining was performed first, following fixation and permeabilization using the Foxp3-staining kit (eBioscience Cat#: 00-5523-00). After staining, all suspensions were washed with FACS buffer, fixed, and analyzed using a LSRII Fortessa or Cytek Aurora Full Spectrum Flow Cytometer. The staining antibodies were used at a dilution of 1:200.

### Western blots

Whole protein extracts were prepared using Cell Lysis Buffer (Cell Signaling Technology, cat. No. 9803). Protein samples were separated by gel electrophoresis using a 4-20% mini-PROTEAN pre-cast gel (BioRad #4561093) with 1X Tris-Glycine-SDS, transferred to a PVDF membrane using the BioRad Trans-Blot SD Semi-Dry Electrophoretic Transfer cell and then incubated with the indicated antibodies after blocking with 5% nonfat milk in TBS-Tween 20 buffer. If necessary, membranes were stripped using OneMinute Western Blot Stripping Buffer (GM Biosciences, Cat. No. GM6001) before re-probing.

### ELISA

Cells were treated with UR238 at the indicated concentrations, and protein samples were generated using Cell Lysis Buffer (Cell Signaling Technology, cat. No. 9803). HE4 levels were determined using the Human HE4/WFDC2 DuoSet ELISA (R&D Systems, cat no. DY6274-05), following the provided protocol. HE4 levels were normalized to total protein concentrations for each sample determined by BioRad DC Protein Assay (Cat. No. 5000111) following the manufacturer’s protocol.

### Statistical analysis

GraphPad Prism 10 software (GraphPad Software Inc.) was used for the generation of all graphs and statistical analyses. P values < 0.05 were determined to be statistically significant. All flow cytometry data was analyzed using FlowJo 10.10.0 software (FlowJo LLC). Error bars in graphs indicate the mean ± SEM. Statistical analysis was performed using Mann-Whitney U tests, unpaired t-tests or Two-Way ANOVA to test for differences between groups.

### Quantitative real-time PCR

RNA collection was performed using the Zymo Research Direct-zol RNA miniprep kit (Cat # R2050) and cDNA was generated using the Bio-Rad iScript cDNA Synthesis Kit (Cat # 1708890). Quantitative PCR was performed in triplicate by loading 2 µl cDNA reaction, 10 µl Power SYBR Green Master mix (Applied Biosystems Cat #: 4368577) or ABI Taqman Master mix (Cat # 4304437), 1 µL of corresponding 20X SYBR primers or Assay-on-demand primers, 7 µL of nuclease-free water. Samples were run on a QuantStudio 12KFlex and data were analyzed using the ΔΔCt method. Relative expression levels were normalized to GAPDH. The primers used were: WFDC2 (Assay ID: Hs00899484_m1, Cat #: 4331182) and GAPDH (Assay ID: Hs02758991_g1, Cat #: 4331182)

### Luminex

Tumor tissue from tumor bearing rats was processed in 500ml of tissue lysis buffer, mechanically homogenized and incubated at room temperature for ∼10 minutes, then placed on ice. Ascites fluid was centrifuged at 14,000 x g for 10 minutes and the cell free supernatant was collected and placed on ice. Protein concentrations were determined using the BioRad DC Protein Assay (Cat. No. 5000111) for normalization. Cytokine analysis was performed using the ProcartaPlex 22-Plex Immunoassay (EPX220-30122-90, Invitrogen) kit according to the manufacturer’s protocol. Samples were run on Bio-Plex 200 system.

## Data availability

The data generated in this study are available in the article or from the corresponding author upon reasonable request.

## Acknowledgements

We thank the Flow Cytometry Core and Genomics Research Core at the University of Rochester for supporting flow cytometry and genomic experiments. We also thank all the patients that donated tissue that was utilized in this work.

## References

1. CDC. Ovarian Cancer Statistics. Ovarian Cancer. 2024 June 12 [accessed 2025 Mar 11]. https://www.cdc.gov/ovarian-cancer/statistics/index.html

2. Kim J, Park E, Kim O, Schilder J, Coffey D, Cho C-H, Bast R. Cell Origins of High-Grade Serous Ovarian Cancer. Cancers. 2018;10(11):433. doi:10.3390/cancers10110433

3. Labidi-Galy SI, Papp E, Hallberg D, Niknafs N, Adleff V, Noe M, Bhattacharya R, Novak M, Jones S, Phallen J, et al. High grade serous ovarian carcinomas originate in the fallopian tube. Nature Communications. 2017;8(1):1093. doi:10.1038/s41467-017-00962-1

4. Matulonis UA, Sood AK, Fallowfield L, Howitt BE, Sehouli J, Karlan BY. Ovarian cancer. Nature Reviews Disease Primers. 2016;2(1):16061. doi:10.1038/nrdp.2016.61

5. Mellman I, Chen DS, Powles T, Turley SJ. The cancer-immunity cycle: Indication, genotype, and immunotype. Immunity. 2023;56(10):2188–2205. doi:10.1016/j.immuni.2023.09.011

6. Zhang L, Conejo-Garcia JR, Katsaros D, Gimotty PA, Massobrio M, Regnani G, Makrigiannakis A, Gray H, Schlienger K, Liebman MN, et al. Intratumoral T Cells, Recurrence, and Survival in Epithelial Ovarian Cancer. New England Journal of Medicine. 2003;348(3):203–213. doi:10.1056/NEJMoa020177

7. Cendrowicz E, Sas Z, Bremer E, Rygiel TP. The Role of Macrophages in Cancer Development and Therapy. Cancers. 2021;13(8):1946. doi:10.3390/cancers13081946

8. Hamanishi J, Mandai M, Ikeda T, Minami M, Kawaguchi A, Murayama T, Kanai M, Mori Y, Matsumoto S, Chikuma S, et al. Safety and Antitumor Activity of Anti–PD-1 Antibody, Nivolumab, in Patients With Platinum-Resistant Ovarian Cancer. Journal of Clinical Oncology. 2015;33(34):4015–4022. doi:10.1200/JCO.2015.62.3397

9. Zhou Q, Liang J, Yang T, Liu J, Li B, Li Y, Fan Z, Wang W, Chen W, Yuan S, et al. Carfilzomib modulates tumor microenvironment to potentiate immune checkpoint therapy for cancer. EMBO Molecular Medicine. 2022;14(1):e14502. doi:10.15252/emmm.202114502

10. Park JE, Miller Z, Jun Y, Lee W, Kim KB. Next-generation proteasome inhibitors for cancer therapy. Translational Research. 2018;198:1–16. doi:10.1016/j.trsl.2018.03.002

11. Manasanch EE, Orlowski RZ. Proteasome inhibitors in cancer therapy. Nature Reviews Clinical Oncology. 2017;14(7):417–433. doi:10.1038/nrclinonc.2016.206

12. Mishto M, Liepe J. Post-Translational Peptide Splicing and T Cell Responses. Trends in Immunology. 2017;38(12):904–915. doi:10.1016/j.it.2017.07.011

13. Almond J, Cohen G. The proteasome: a novel target for cancer chemotherapy. Leukemia. 2002;16(4):433– 443. doi:10.1038/sj.leu.2402417

14. Rajkumar SV, Richardson PG, Hideshima T, Anderson KC. Proteasome Inhibition As a Novel Therapeutic Target in Human Cancer. Journal of Clinical Oncology. 2005;23(3):630–639. doi:10.1200/JCO.2005.11.030

15. Harrington BS, Annunziata CM. NF-κB Signaling in Ovarian Cancer. Cancers. 2019;11(8):1182. doi:10.3390/cancers11081182

16. Ramirez PT, Landen CN, Coleman RL, Milam MR, Levenback C, Johnston TA, Gershenson DM. Phase I trial of the proteasome inhibitor bortezomib in combination with carboplatin in patients with platinum- and taxane-resistant ovarian cancer. Gynecologic Oncology. 2008;108(1):68–71. doi:10.1016/j.ygyno.2007.08.071

17. Aghajanian C, Dizon DS, Sabbatini P, Raizer JJ, Dupont J, Spriggs DR. Phase I Trial of Bortezomib and Carboplatin in Recurrent Ovarian or Primary Peritoneal Cancer. Journal of Clinical Oncology. 2005;23(25):5943–5949. doi:10.1200/jco.2005.16.006

18. Troy Pellom S. Development of Proteasome Inhibitors as Therapeutic Drugs. Journal of Clinical & Cellular Immunology. 2012 [accessed 2025 July 10];01(S5). https://www.omicsonline.org/development-of-proteasome-inhibitors-as-therapeutic-drugs-2155-9899.S5-005.php?aid=5237. doi:10.4172/2155-9899.s5-005

19. Redic K. Carfilzomib: a novel agent for multiple myeloma. Journal of Pharmacy and Pharmacology. 2013;65(8):1095–1106. doi:10.1111/jphp.12072

20. Field-Smith A, Morgan GJ, Davies FE. Bortezomib (Velcade) in the treatment of multiple myeloma. Therapeutics and Clinical Risk Management. 2006;2(3):271–279. doi:10.2147/tcrm.2006.2.3.271

21. Maniati E, Berlato C, Gopinathan G, Heath O, Kotantaki P, Lakhani A, McDermott J, Pegrum C, Delaine-Smith RM, Pearce OMT, et al. Mouse Ovarian Cancer Models Recapitulate the Human Tumor Microenvironment and Patient Response to Treatment. Cell Reports. 2020;30(2):525–540.e7. doi:10.1016/j.celrep.2019.12.034

22. Soong TR, Howitt BE, Horowitz N, Nucci MR, Crum CP. The fallopian tube, “precursor escape” and narrowing the knowledge gap to the origins of high-grade serous carcinoma. Gynecologic Oncology. 2019;152(2):426–433. doi:10.1016/j.ygyno.2018.11.033

23. Meza-Perez S, Randall TD. Immunological Functions of the Omentum. Trends in Immunology. 2017;38(7):526–536. doi:10.1016/j.it.2017.03.002

24. Chen X, Li Y, Xia H, Chen YH. Monocytes in Tumorigenesis and Tumor Immunotherapy. Cells. 2023;12(13):1673. doi:10.3390/cells12131673

25. Liu M, Silva-Sanchez A, Randall TD, Meza-Perez S. Specialized immune responses in the peritoneal cavity and omentum. Journal of Leukocyte Biology. 2021;109(4):717–729. doi:10.1002/JLB.5MIR0720-271RR

26. Li Y, Zhang Y, Pan G, Xiang L, Luo D, Shao J. Occurrences and Functions of Ly6Chi and Ly6Clo Macrophages in Health and Disease. Frontiers in Immunology. 2022;13:901672. doi:10.3389/fimmu.2022.901672

27. Wang S, Wang J, Chen Z, Luo J, Guo W, Sun L, Lin L. Targeting M2-like tumor-associated macrophages is a potential therapeutic approach to overcome antitumor drug resistance. npj Precision Oncology. 2024 [accessed 2025 July 8];8(1). https://www.nature.com/articles/s41698-024-00522-z. doi:10.1038/s41698-024-00522-z

28. Hellstrom I, Raycraft J, Hayden-Ledbetter M, Ledbetter JA, Schummer M, McIntosh M, Drescher C, Urban N, Hellstrom KE. The HE4 (WFDC2) Protein Is a Biomarker for Ovarian Carcinoma.

29. Moore RG, Hill EK, Horan T, Yano N, Kim K, MacLaughlan S, Lambert-Messerlian G, Tseng YD, Padbury JF, Miller MC, et al. HE4 (WFDC2) gene overexpression promotes ovarian tumor growth. Scientific Reports. 2014;4(1):3574. doi:10.1038/srep03574

30. Moore RG, McMeekin DS, Brown AK, DiSilvestro P, Miller MC, Allard WJ, Gajewski W, Kurman R, Bast RC, Skates SJ. A novel multiple marker bioassay utilizing HE4 and CA125 for the prediction of ovarian cancer in patients with a pelvic mass. Gynecologic Oncology. 2009;112(1):40–46. doi:10.1016/j.ygyno.2008.08.031

31. Rowswell-Turner RB, Singh RK, Urh A, Yano N, Kim KK, Khazan N, Pandita R, Sivagnanalingam U, Hovanesian V, James NE, et al. HE4 Overexpression by Ovarian Cancer Promotes a Suppressive Tumor Immune Microenvironment and Enhanced Tumor and Macrophage PD-L1 Expression. The Journal of Immunology. 2021;206(10):2478–2488. doi:10.4049/jimmunol.2000281

32. Huang Z, Wu Y, Zhou X, Xu J, Zhu W, Shu Y, Liu P. Efficacy of Therapy with Bortezomib in Solid Tumors: A Review based on 32 Clinical Trials. Future Oncology. 2014;10(10):1795–1807. doi:10.2217/fon.14.30

33. Dimopoulos MA, Goldschmidt H, Niesvizky R, Joshua D, Chng W-J, Oriol A, Orlowski RZ, Ludwig H, Facon T, Hajek R, et al. Carfilzomib or bortezomib in relapsed or refractory multiple myeloma (ENDEAVOR): an interim overall survival analysis of an open-label, randomised, phase 3 trial. The Lancet Oncology. 2017;18(10):1327–1337. doi:10.1016/s1470-2045(17)30578-8

34. Liu R, Lu S, Deng Y, Yang S, He S, Cai J, Qiang F, Chen C, Zhang W, Zhao S, et al. PSMB4 expression associates with epithelial ovarian cancer growth and poor prognosis. Archives of Gynecology and Obstetrics. 2016;293(6):1297–1307. doi:10.1007/s00404-015-3904-x

35. Brüning A, Burger P, Vogel M, Rahmeh M, Friese K, Lenhard M, Burges A. Bortezomib treatment of ovarian cancer cells mediates endoplasmic reticulum stress, cell cycle arrest, and apoptosis. Investigational New Drugs. 2009;27(6):543–551. doi:10.1007/s10637-008-9206-4

36. Lee YJ, Seol A, Lee M, Kim J-W, Kim HS, Kim K, Suh DH, Kim S, Kim SW, Lee J-Y. A Phase II Trial to Evaluate the Efficacy of Bortezomib and Liposomal Doxorubicin in Patients With BRCA Wild-type Platinum-resistant Recurrent Ovarian Cancer (KGOG 3044/EBLIN). In Vivo. 2022;36(4):1949–1958. doi:10.21873/invivo.12917

37. El-Arabey AA, Alkhalil SS, Al-Shouli ST, Awadalla ME, Alhamdi HW, Almanaa TN, Mohamed SSEM, Abdalla M. Revisiting macrophages in ovarian cancer microenvironment: development, function and interaction. Medical Oncology. 2023;40(5):142. doi:10.1007/s12032-023-01987-x

38. Ruytinx P, Proost P, Van Damme J, Struyf S. Chemokine-Induced Macrophage Polarization in Inflammatory Conditions. Frontiers in Immunology. 2018 [accessed 2025 July 8];9. https://www.frontiersin.org/article/10.3389/fimmu.2018.01930/full. doi:10.3389/fimmu.2018.01930

39. Bouchard D, Morisset D, Bourbonnais Y, Tremblay GM. Proteins with whey-acidic-protein motifs and cancer. The Lancet Oncology. 2006;7(2):167–174. doi:10.1016/S1470-2045(06)70579-4

40. Kim K, Khazan N, McDowell JL, Snyder CWA, Miller JP, Singh RK, Whittum ME, Turner R, Moore RG. The NF-κB-HE4 axis: A novel regulator of HE4 secretion in ovarian cancer Tan W, editor. PLOS ONE. 2024;19(12):e0314564. doi:10.1371/journal.pone.0314564

41. Pakjoo M, Ahmadi SE, Zahedi M, Jaafari N, Khademi R, Amini A, Safa M. Interplay between proteasome inhibitors and NF-κB pathway in leukemia and lymphoma: a comprehensive review on challenges ahead of proteasome inhibitors. Cell Communication and Signaling. 2024;22(1):105. doi:10.1186/s12964-023-01433-5

