## Supplemental Figure 1 for "Effects of a novel proteasome inhibitor, UR238 on the tumor immune microenvironment and growth in epithelial ovarian cancer"

**Supplemental Figures**

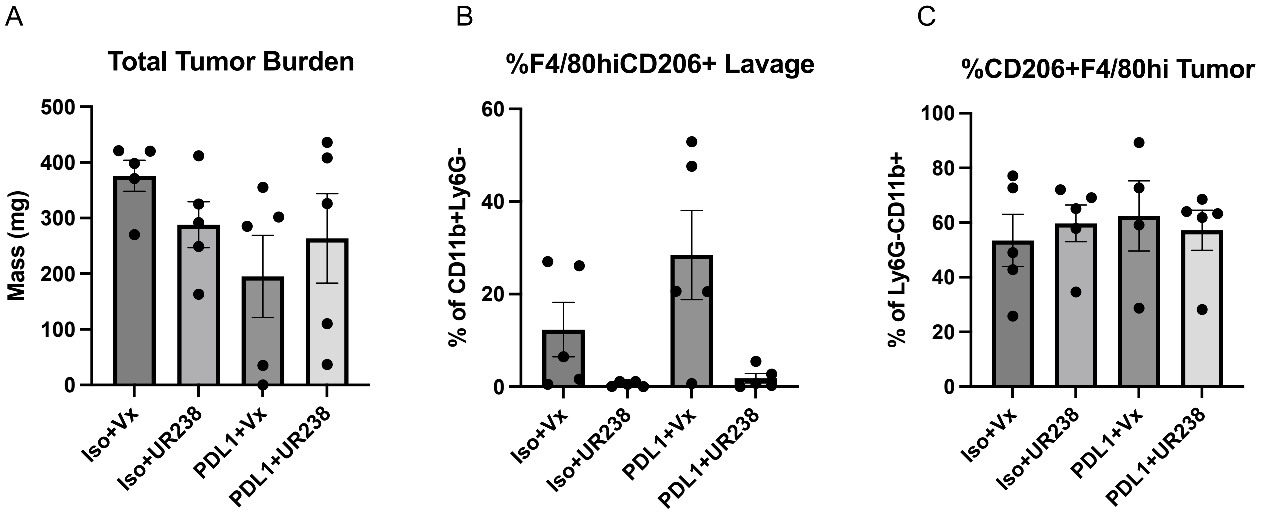


**Figure S1: UR238 + anti-PD-L1 does not synergize to improve response in HGS3 tumors.**

A) Total tumor burden of HGS3 tumors in the indicated treatment group. B) Percentage of F4/80$^{hi}$CD206$^{+}$ M2 macrophages in the peritoneal cavity in the indicated treatment groups. C) Percentage of CD206$^{+}$F4/80$^{hi}$ within HGS3 tumors in the indicated treatment groups.
